# Long-term molecular evolutionary rate determines intraspecific genetic diversity

**DOI:** 10.1101/846824

**Authors:** Jiaqi Wu, Takahiro Yonezawa, Hirohisa Kishino

**Affiliations:** School of Life Science and Technology, Tokyo Institute of Technology, Meguro Ward, Tokyo 152-8550, Japan; Faculty of Agriculture, Tokyo University of Agriculture, Atsugi City, Kanagawa 243-0034, Japan; Graduate School of Agricultural and Life Sciences, The University of Tokyo, Bunkyo Ward, Tokyo 113-8657, Japan

## Abstract

What determines genetic diversity and how it connects to the various biological traits is unknown. In this work, we offer answers to these questions. By comparing genetic variation of 14,671 mammalian gene trees with thousands of individual genomes of human, chimpanzee, gorilla, mouse and dog/wolf, we found that intraspecific genetic diversity is determined by long-term molecular evolutionary rates, rather than *de novo* mutation rates. This relationship was established during the early stage of mammalian evolution. Expanding this new finding, we developed a method to detect fluctuations of species-specific selection on genes as the deviations of intra-species genetic diversity predicted from long-term rates. We show that the evolution of epithelial cells, rather than of connective tissue, mainly contributes to morphological evolution of different species. For humans, evolution of the immune system and selective sweeps subjected by infectious diseases are most representative of adaptive evolution.

## Introduction

Genomes vary among species and individuals within a single species. What determines the genetic polymorphisms is one of the core questions in evolutionary biology. Mutation is the source of both intra- and interspecific genetic diversity. Within a species, each new mutation may form a new polymorphic site. If a mutation is finally fixed in the species, it becomes a substitution and contributes to the interspecific genetic diversity. Source of mutations is replication errors in genomes in germline cells, the so-called *de novo* mutations. The genome-wide distribution of *de novo* mutations may be related to recombination hotspots [1]. However, such a relationship cannot explain the patterns of genome-wide distribution of polymorphic sites. In addition, although nucleotide diversity is generally regarded as being determined by the effective population size (*N*_*e*_), Lewontin’s paradox tells us that distribution of diversity levels and variation in population size differ by many orders of magnitude, and the former is much narrower [2]. Furthermore, the causality of intra- and interspecific genetic diversity is also under debate. Because mutations occur in individuals, many people believe that intraspecific genetic diversity determines interspecific diversity. However, little is known about the connections between individual-level random mutation and species-level substitutions.

As pointed out by R. A. Fisher, the majority of mutations in a population are deleted by chance [3]. According to Motoo Kimura’s neutral theory of molecular evolution, most mutations are either deleterious or selectively neutral [4, 5]. Hence, the molecular evolutionary rate of a gene is the product of the mutation rate and the proportion of neutral mutations [6]. In contrast, intraspecies genetic diversity is generated by mutations occurring along genealogies in a population that have neither reached fixation nor been lost. The shape of a genealogy depends on population history and effective population size, and the lifespan of a mutation depends additionally on the selection intensity [7, 8]. Here, we note that all genes in a genome share the same demographic history. Among-gene variation in genetic diversity thus largely reflects among-gene variation in the proportion of neutral mutations, as lineages originating from deleterious mutants are too short-lived to substantially contribute to present genetic diversity. Because among-gene variation in molecular evolutionary rates is also determined by the among-gene variation in a proportion of neutral mutations [9], we arrive at the hypothesis that long-term molecular evolutionary rates affect intraspecies genetic diversity.

If our hypothesis is correct, the genetic diversity within species can be predicted from the long-term molecular evolutionary rates. Based on this idea, a novel genome evolutionary method to detect the signals of species-specific evolution becomes possible as well, because the residues of the observed and predicted genetic diversities can be regarded as the signals of species-specific selection. Ever since Darwin, species-specific evolution is regarded as a key factor for the diversification of living organisms and their remarkable adaptations to their environments. Nevertheless, the genetic background of species-specific evolution, and especially how selection pressures fluctuate along the history of species, is still unclear even in this genomic era. This difficulty largely stems from our limited knowledge about what determines genetic diversity and how genetic polymorphism is shaped within species.

The aims of this study are (1) to clarify how genetic polymorphisms have been shaped within species by testing our hypothesis that long-term molecular evolutionary rates affect intraspecies genetic diversity, based on phylogenomic analysis of mammals and population genomic analysis of five species (human, chimpanzee, gorilla, mouse and dog/wolf), and (2) to develop a new genome evolutionary method to detect the signals of species-specific selection. We proved that long-term molecular evolutionary rates determine intraspecific genetic diversity, and reported species-specific evolution of human, chimpanzee, gorilla, mouse and dog/wolf. We demonstrated that the evolution of epithelial cells, rather than connective tissue, mainly contributes to the morphological evolution of different species. Selection pressures operating on the gene group related to immune and infectious disease have been strengthened specifically in human. This new method opens new windows to elucidate species-specific evolution.

## Results and Discussions

### Measurement of genetic polymorphisms and *de novo* mutations

To test the above hypothesis, that long-term molecular evolutionary rates affect intraspecies genetic diversity, we analyzed and compared the genetic diversity of humans with the mammalian molecular evolutionary rates of each gene, focusing on protein-coding regions. The different evolutionary time scales of mammals and humans, including examples of gene trees and intraspecies genetic diversity, are shown in Figure S1. Indices of genetic diversity capture different age groups of surviving mutations. Depending on their unique population histories, the effect of long-term molecular evolutionary rates on intraspecies genetic diversity may vary among species. The proportion of segregating sites (*q*) is the most direct estimate of human intraspecies genetic diversity. Nucleotide diversity (π), the mean pairwise distance among sequences, more heavily weights older mutations. Singletons are the collection of recent mutations along the terminal edges of a genealogy. The allele frequency of singletons in autosomes is 1/5,008, which is less than 0.02% of the number of sites in 2,504 humans in the 1000 Genomes Project data [10-12]. *De novo* mutations are obtained by comparing genomes of direct offspring with those of their parents, with the differences uncovered being the closest reflection of germ cell mutations [1]. By comparing the effects of molecular evolutionary rates on these different indices, we may gain some sense of the longevity of deleterious or slightly deleterious mutations that have been selected against [5].

In each population, natural selection works on phenotypes as well as on the sets of genes that control them. Depending on the surrounding environment, the targets of natural selection may vary among species. To identify species-specific targets, we searched for significant deviations from the genetic diversity predicted by long-term molecular evolutionary rates. Genes with an unexpectedly high genetic diversity may be under reduced functional constraints, whereas those with an unexpectedly low genetic diversity may be subject to enhanced constraints.

### Bridging micro- and macroevolution at the molecular level

We obtained 14, 671 genes across 96 mammalian species, ranging from platypus to human, and calculated median branch lengths of nucleotide gene trees as an estimate of among-gene variation in mammalian molecular evolutionary rates. Because such rate variation has arisen during 185 million years of mammalian evolution [13], we refer to these rates as long-term rates. In addition, we collected genomes of 2,504 humans from the 1000 Genomes Project (phase 3 data [10-12]) and calculated the proportion of segregating sites (*q*) as the among-gene variation in human genetic polymorphism (Figure 1A). The time to the most recent common ancestor of modern humans (*Homo sapiens*) is only 202,000 years [14], which corresponds to approximately 0.1% of the mammalian time scale (Figure S1). However, as expected, strong linear relationships were found between mammalian long-term rates and human *q* (Figure 1A, cor = 0.585, *p* < 2 × 10^−16^).

**Figure 1.**
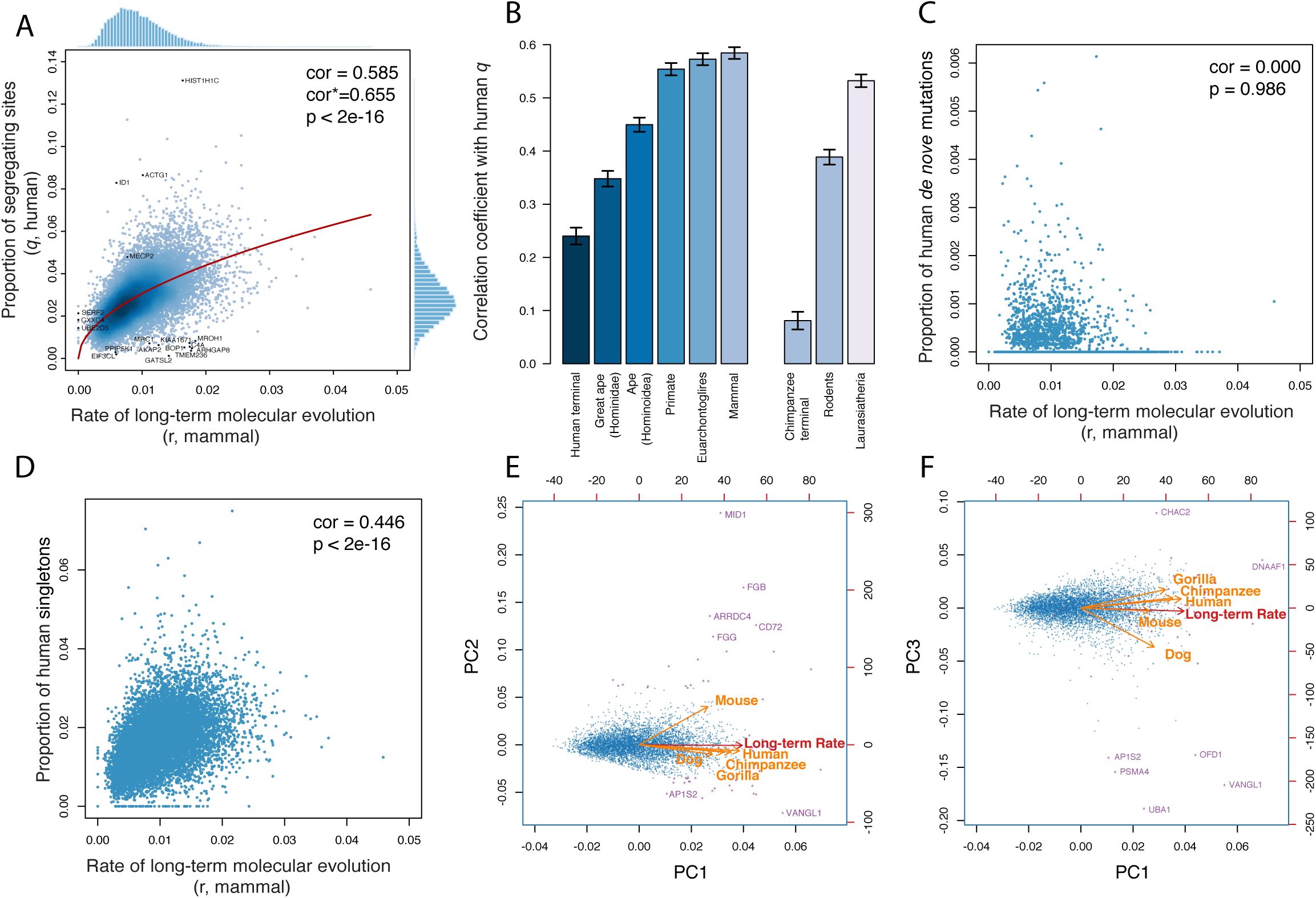
Bridging micro- and macroevolution at the molecular level. **A**. Correlation between the human proportion of segregating sites (*q*) and the rate of mammalian molecular evolution (*r*) of 14, 671 genes. **B**. Correlation between the human proportion of segregating sites (*q*) and the rate of molecular evolution (*r*) based on gene trees of different taxonomic clades. **C**. Correlation between the proportion of human *de novo* mutations and the rate of mammalian molecular evolution (*r*). **D**. Correlation between the proportion of human singletons and the rate of mammalian molecular evolution (*r*). **E-F**. Principal component analysis (PCA) of *q* of five species and the long-term rate using 5,560 single-copy genes.

Because uncertainties in branch lengths, the randomness of mutations, and the stochasticity of genealogies deflate the values of net correlations, differences among species in these correlations are difficult to evaluate (see Methods). Here, we note that the variance in the proportion of segregating sites (*q*) generally takes the form of the sum of two terms. The first term reflects random mutations and can be assumed to follow a Poisson distribution. The second term, the stochasticity of genealogies, is proportional to the square of the mean [8, 15]. This variance formula is consistent with that of a negative binomial distribution, which takes into account over-dispersion relative to the Poisson distribution. Because the coefficient of the second term is determined by factors such as demographic histories, it can be assumed to be constant among genes. We therefore conducted a negative binomial regression of human *q* on mammalian long-term rates, with gene length included as an offset (Figure 1A, red curve). The variances in human *q* were obtained using the estimated shape parameter of the negative binomial distribution. With the estimated variances, we corrected the correlation between mammalian long-term rates and human *q*. The bias-corrected correlation was as high as 0.665 (Figure 1A, *p* < 2 × 10^−16^), which indicates that the intraspecies genetic diversity of a gene is thus largely influenced by the long-term evolutionary rate of the gene. Variation in functional constraints on genes is the likely source of this causal relation [16]. The estimated shape parameter in the negative binomial distribution in human was as large as 125,785, thus indicating that over-dispersion of human *q* can be ignored, and human *q* largely followed a Poisson distribution. We tested four non-human mammals, including chimpanzee, orangutan, mouse and dog/wolf. The correlation between *q* of these 4 species and the long-term rates ranged from 0.368 (mouse) to 0.478 (chimpanzee), with is consistent with the findings in human (Figure S2, all with *p* < 2 × 10^−16^).

### Duration of the bridge between micro- and macroevolution

Functional constraints vary among lineages [9], and molecular evolutionary rates accordingly vary as well. Consequently, intraspecies genetic diversity may correlate more strongly with molecular evolutionary rates in subtrees of closely related species. The size of the most influential subtree may depend on the scale of variability. We thus also wished to address the question of how long the “long-term” molecular evolutionary rate of a gene affects intraspecies diversity.

Figure 1B shows the raw correlation between the human proportion of segregating sites (*q*) and molecular evolutionary rate variation inferred from different taxonomic hierarchies, ranging from the human/chimpanzee divergence (human terminal branches) to family Hominidae (great apes) to superorder Euarchontoglires. Significant correlations between human *q* and rate variations were found for all clades, with higher taxonomic hierarchies showing stronger correlations. Furthermore, among-gene rate variation of rodents and Laurasiatheria correlated highly with human *q* (cor = 0.389 and 0.532, respectively; both *p* < 2 × 10^−16^). The rate variation of mammal had the highest raw correlation with human *q* (Figure 1B, cor = 0.585). Taken together, these results indicate that long-term molecular evolutionary rates were established at the early stage of the origin of mammals, thus acting as the main force impacting the intraspecies genetic diversity of mammalian species.

### The immediate effect of natural selection

Among-gene variation of intraspecies genetic diversity is caused by among-gene variation of mutation rate and among-gene variation of functional constraints. To examine whether the long-term molecular evolutionary rate of a gene correlates with its mutation rate, we regressed the rate of point mutations per generation (proportion of human *de novo* mutations) on mammalian long-term rates. We found no correlation between the proportion of *de novo* mutations and mammalian long-term rates (cor = 0.00, *p* =0.986, Figure 2C). In contrast, the proportion of singletons in the 1000 Genomes Project human data was found to be affected by mammalian long-term rates (cor = 0.446, *p* < 2 × 10^−16^; Figure 2D). The expected number of singletons was estimated as

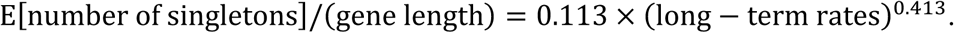

**Figure 2.**
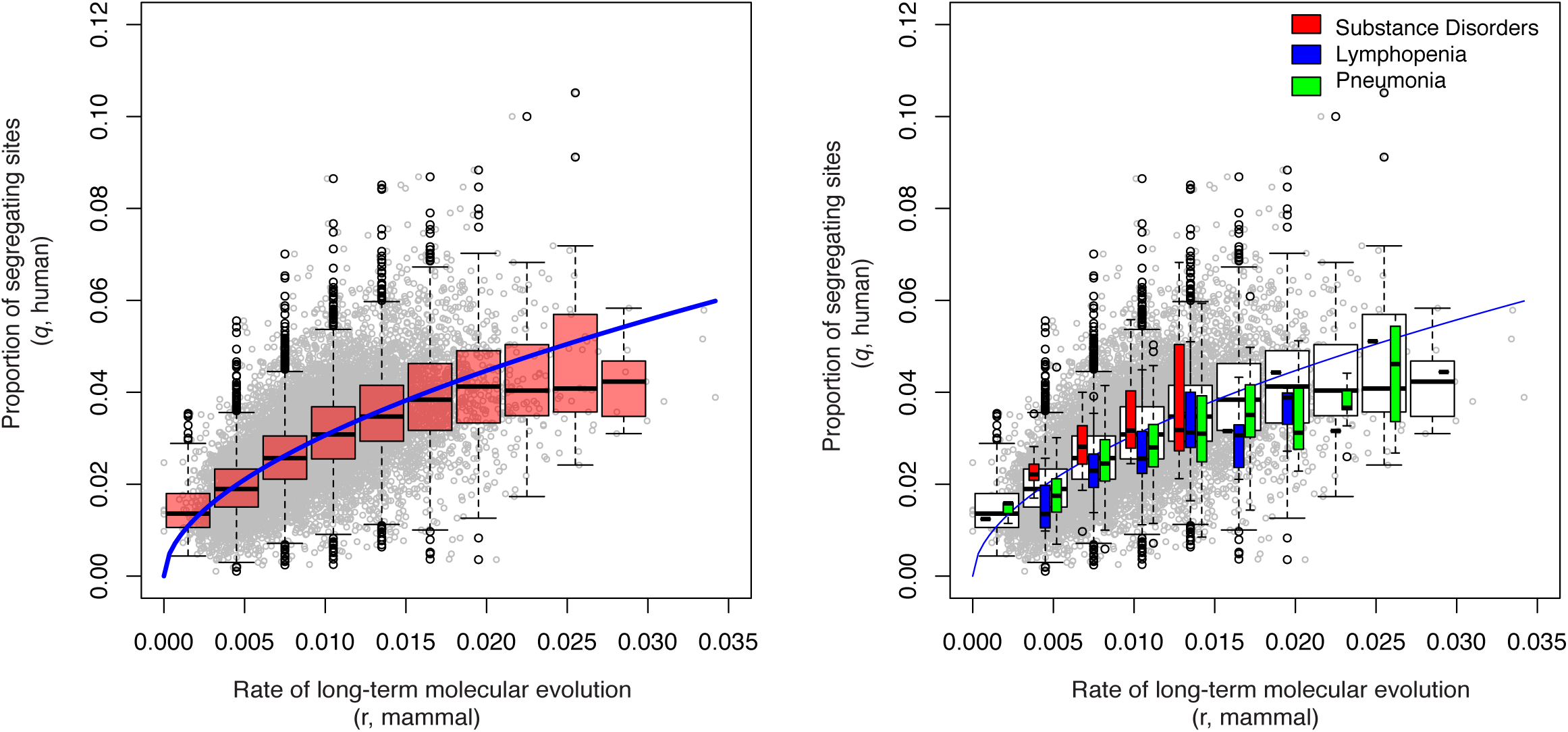
Method of disease-associated gene set test. **A**. Scatter plot, boxplot and regression model of the human proportion of segregating sites (*q*) vs. mammalian molecular evolutionary rates of 14,671 genes. **B**. Examples of significant disease-associated gene sets in humans revealed by analysis of 14,267 disease-associated gene sets. Genes related to the following diseases in humans are shown as examples: substance-related disorders, lymphopenia and pneumonia.

The effect of long-term molecular evolutionary rate was highly significant (*p* < 2 × 10^−16^, Table 1).

**Table 1.** Negative binomial regression of numbers of human *de novo* mutations and singletons on the median branch length of the mammalian gene trees. The regression model is logsE[counts]/(gene length)t = *α* + *β* log(med*i*an branch length), that is, E[counts]/(gene length) = *e*^α^(med*i*an branch length)^β^.

|  | *de novo* mutation |  |  | singleton |  |  |
| --- | --- | --- | --- | --- | --- | --- |
|  | estimate | SE | p-value | estimate | SE | p-value |
| intercept | -9.938 | 0.307 | $<2*10^{-16}$ | -2.177 | 0.026 | $<2*10^{-16}$ |
| log(median_BL) | -0.008 | 0.064 | 0.904 | 0.413 | 0.006 | $<2*10^{-16}$ |

Singletons, which reflect relatively recent changes, may help shape complex traits [17, 18]. Selection, caused by long-term evolutionary force, has in turn influenced human genetic diversity over a short time scale, probably soon after the occurrence of the initial mutation.

### Commonality and specificity among species

The among-genes variation of long-term molecular evolutionary rate reflects the variable strength of functional constraints, which in turn largely determines the intra-species genetic diversity. To visualize the commonality and specificity between species, we conducted principal component analysis (PCA) (Figures 1E-F). The first principal component (PC1) explains 48% of variability and expresses the size of inter- and intraspecies genetic diversity as a whole. PC2 has a contribution of 13.3% and represents reduced/enhanced functional constraints in mouse. PC3 has a contribution of 13.0% and expresses enhanced/reduced functional constraints in dog/wolf. The relationships among the five species precisely reflect their phylogenetic relationships.

To interpret the genes that comprise the principal components, we examined the functions of the top 20 genes for each of the three PCs (Table S1; for enrichment search [19], we set FDR (false discovery rate) 0.05 as the significance cut-off value). The top 20 genes with negative deviations in PC1 are generally those genes that have enhanced functional constraints in mammals. These genes are enriched for the categories of learning or memory (GO:0007611, FDR = 0.0035) and behavior (GO:0007610, FDR = 0.004) (Table S1) in a Biological Process (GO) enrichment search [20]. The top 20 genes in the positive direction of PC2, which have relaxed functional constraints or have been subjected to diversifying selection in mouse, are enriched for fibrinogen complex (GO:0005577, FDR = 1.87E-05, Table S1). The MID1 gene, which has the largest deviation in PC2, is previously reported to have undergone X-inactivation in human, while escaping in mouse [21]. For the top 20 genes in PC3, a GO enrichment search did not suggest any significant categories. However, a Reactome Pathways search [22] found that these genes are enriched mostly for Class I MHC-mediated antigen processing & presentation and the adaptive immune system (seven genes in 20 (Table S1): AP1S2, CDC27, PSMA4, PSMC2, TLR4, UBA1, UBE2L6). The adaptive immune system is subjected to enhanced functional constraints in dog/wolf, probably because its carnivorous lifestyle in the wild ancestors was accompanied by a higher risk of pathogen infections.

### Species-specific evolution of genes

Species-specific enhanced/reduced functional constraints of each gene can be quantified by measuring the deviation in the proportion of segregating sites (*q*) from predictions based on long-term molecular evolutionary rates. We standardized the deviation by transforming the negative binomial regression to a standard normal distribution. First, we obtained the lower tail *p*-value of a gene by contrasting the observed *q*-value with a negative binomial distribution based on the predicted value and the estimated shape parameter. This *p*-value was mapped to the quantile of the standard normal distribution. A positive value for the deviation indicates a reduction in functional constraints or the presence of diversifying selection, whereas a negative value reflects enhancement of functional constraints or the occurrence of purifying selection (Figure 2).

We obtained negative log-transformed *p*-values for each gene, and conducted correspondence analysis (CA) of the five species, to find the genes that could explain each species mostly. The results are summarized in Figure 3A. With FDR set to 0.01, we detected 53 genes whose diversity can explain the species-specific diversities, among which 24 were designated as being related to disease by the UniProt database [23] (Table S2). Based on their locations, we clustered the genes into three groups: ape-group, mouse-group and dog/wolf-group. Mouse-group is enriched for structural molecule activity (GO: 0005198, FDR = 0.003), while the other two groups do not have any significant enrichment. Notably, the genes detected by PCA and correspondence analysis (CA) largely overlap, especially those that are most special for different PCs and groups (e.g., MID1 and FGB for mouse, UBA1 and VANGL1 for dog/wolf, Figure 1E-F, 3A).

**Figure 3.**
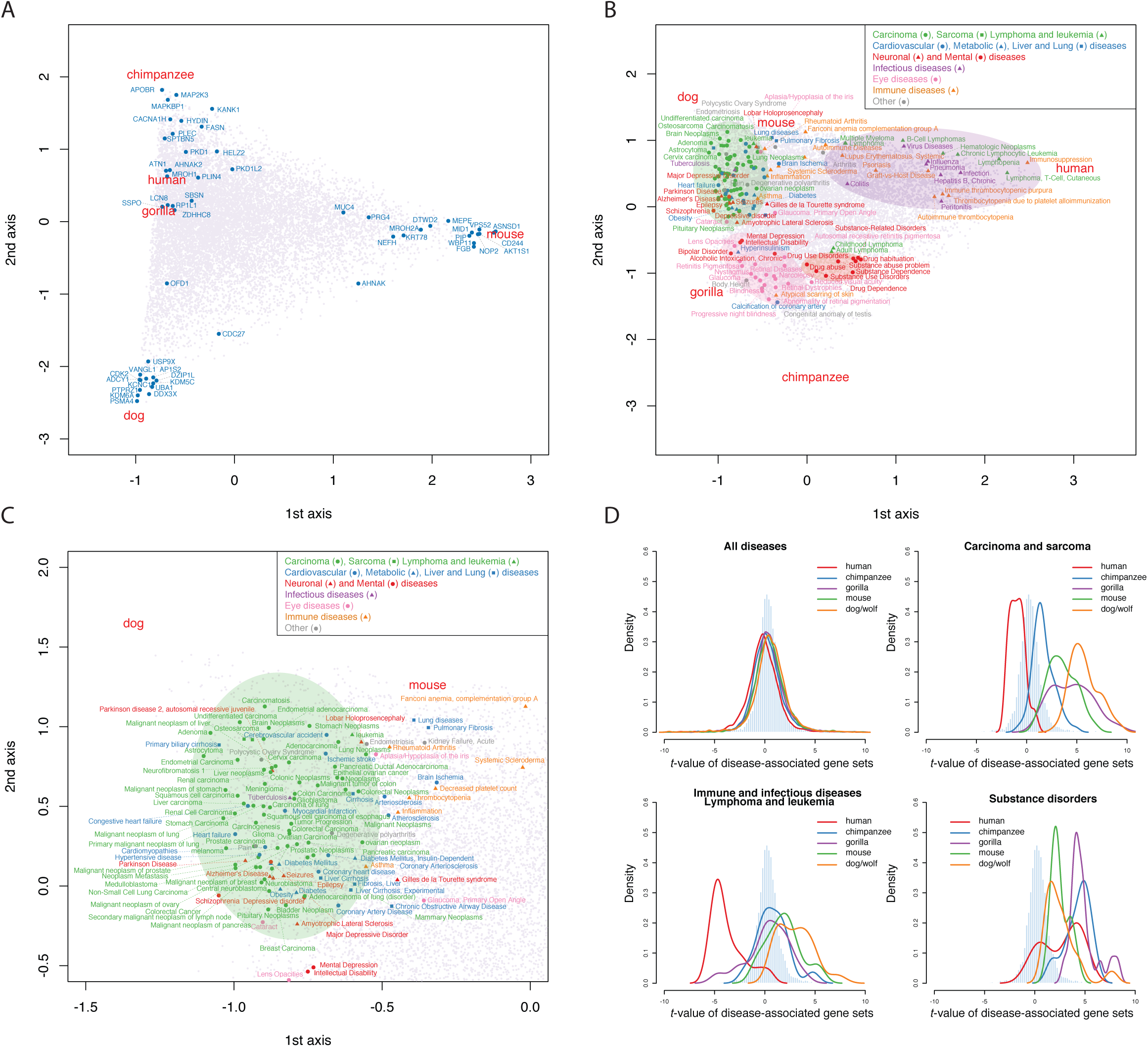
Correspondence analysis on gene and disease-associated gene sets. **A**. Correspondence analysis on 14, 671 genes of five species using –*log*_10_(*p*) of each gene. **B-C**. Correspondence analysis on 14,267 disease-associated gene sets of five species using –*log*_10_(*p*) of each disease-associated gene set. The diseases highlighted are significant diseases with a false discovery rate (FDR) of 0.01. **D**. Distribution of *t*-values of all diseases, as well as significant disease-associated gene sets detected by correspondence analysis. The pale blue histogram indicates the distribution of average t-values of five species of each disease.

### Multiple species disease-associated gene set test

To further examine species-specific constraints on traits, we analyzed the average deviation of disease-associated gene sets. In doing so, we hoped to identify the species specificity of biological traits that are responsible for various diseases. We first downloaded disease-associated gene sets from the DisGeNET database [24]. To obtain a robust result based on multiple genes, we focused on gene sets that included at least two genes contained in our own dataset. Consequently, we analyzed 14,267 disease-related gene sets. To obtain the average deviation of a gene set from predicted genetic diversities, we converted the *p*-values of genes in the gene set to quantiles of the standard normal distribution. The average deviation was measured as the *t*-value of the quantiles (Figure 2, Table S3; see Methods). Using an FDR of 0.01, we identified 20 disease-associated gene sets for humans, 23 for chimpanzee, 86 for gorilla, 31 for mouse and 130 for dog/wolf (Table S4). Even though the variation detected in a single gene may be minor, the effect of a set of genes might show a clear sign of species-specific adaptation.

To capture the patterns of species-specific evolution on the traits revealed by disease-associated genes, we conducted CA based on the negative log-transformed *p*-values (Figure 3B-C). With an FDR of 0.01, 180 gene sets were significant for at least one species (Table S5). These can be assigned to six major categories including cancer diseases (such as carcinoma, sarcoma, lymphoma and leukemia), genetic diseases (such as cardiovascular, metabolic, liver and lung diseases), neuronal and mental diseases, infectious diseases, eye diseases and immune diseases based on the disease category annotations in DisGenNET [24] and the MalaCard database [25]. We marked particular clustering of disease-associated gene sets near each species, and defined 4 four groups: “cancer and organ diseases” group, “immune and infectious diseases” group, “substance disorders” group and “eye diseases” group. Figure 3D shows the distributions of *t*-values of major diseases in each group. Notably, the significant diseases detected for each species (Table S4) and CA (Table S5) are also largely consistent.

### Cancer disease gene sets and morphological evolution

The prominent diseases detected for mouse and dog/wolf are largely cancer diseases with positive *t*-value (Figure 3D, Table S4). The “cancer and organ diseases group” also locates near mouse and dog/wolf in CA analysis (Figure 3B-C). These results indicate that the development of organs is the most significant character that differentiated mouse and dog/wolf from great apes.

Carcinomas are epithelial cell-derived cancers, while sarcomas are connective tissue-derived cancers. Our testing datasets included both types of cancers, but the prominent cancer diseases detected for each species by CA analysis are largely carcinoma of different organs. Only one sarcoma (osteosarcoma) was detected. The evolution of epithelial cells rather than connective tissue may thus be the main contributor to the morphological evolution of different species.

Interestingly, great apes, especially human, have relatively low t-values of cancer diseases compared with mouse and dog/wolf (Figure 3D). Compared with those species, cancer-related genes in great apes are subjected to enhanced functional constraints. Humans have evolved much longer lifespans than other great apes, while dog/wolf and mouse live for fewer years [26, 27]. Cancer-associated gene sets that are subjected to enhanced functional constraints in human may be related to the evolutionary acquisition of longer lifespan.

### Echoes of infectious diseases in the human genome

Outliers in human are largely infectious diseases, immune diseases, lymphoma and leukemia, with negative *t*-values (FDR 0.01). The “immune and infectious disease” group also locates in human in CA (Figure 3B). Influenza-related gene sets have the highest significance in human, with *t*-value -5.62, as well as pneumonia (*t*-value = -5.17) and chronic hepatitis B (*t*-value = -4.51). With FDR as 0.1, more infectious disease-related gene sets were detected in human, such as Saint Louis encephalitis, adult T-cell lymphoma/leukemia, hepatitis C, dengue fever and brucellosis. Several of these diseases have caused multiple historically important pandemics and taken millions of lives (e.g., Saint Louis encephalitis and dengue fever), while some of them still cause severe public health problems now (e.g., influenza and pneumonia). These diseases affected human populations worldwide, leading to a strong selective sweep. The origin and development of agriculture results the population expansions, civilizations, collision of cultures, and globalization. These events might make the exposure risk of the infection disease higher.

### Social structure and natural reward pathway in great apes

Human, chimpanzee and gorilla shared several outliers of substance-related disorders, and they are all highly significant (Table S4). In CA, a related group also locates in the middle of three great apes. Substance-related disorders are mainly defined by a reliance and/or abuse of a certain substance. The mechanism underlying these disorders stems from the effect of the “substance” (e.g., a drug) on the natural reward pathway of the motivational-emotional system. In nature, reward pathways generate positive emotions that motivate organisms to search for food, water, mates and other beneficial environmental resources, and to avoid predators [28]. Great apes display much more complex social behavior compared with all other mammals; in particular, humans have the most complex societal culture of all living organisms and enjoy the largest array of “natural rewards”, including science, art, literature and music. Motivation is a basic driver of human behavior, and positive emotion is at its core [29]. The positive *t*-values associated with substance-related disorders imply that their specificity among mammals is due to evolution of the natural reward pathway in the motivational-emotional system.

We also note, in the vicinity of great apes, several gene sets associated with diseases affecting retina development and eye structures. This may be related to the evolution of color vision in primates: in eutherian mammals, only primates have true trichromatic color vision [30].

## Conclusion

In this work, we provide new insight to understand how polymorphisms are shaped. The distribution of intraspecific polymorphism sites is determined by long-term molecular evolutionary rates, rather than *de novo* mutation rates. The population genomic data of human, chimpanzee, gorilla, mouse and dog/wolf consistently indicated that the impact of effective population size and demographic history of each species on intraspecific genetic diversity is limited. New mutations were subjected to natural selection soon after their emergence, and a large quantity of *de novo* mutations actually does not contribute to intraspecific genetic diversity. The intraspecific genetic polymorphism rapidly approximates to the genetic diversity determined by the long-term molecular evolutionary rates. Such a relationship became established in the early stage of mammalian evolution. The impact of natural selection on shaping intraspecific genetic diversity is much stronger than we expected previously. These findings are different from traditional population genetics, which considers that both the occurrence of mutations and their fixation process are largely random. The chance of survival of a mutation differs depending on the gene locus in which it occurs. Although we did not analyze non-coding regions, we would expect them also to show a similar tendency. At the genetic level, evolution from early mammals to modern humans is indeed a continuous process.

## Materials and Methods

### The probability of intraspecies polymorphism and measurement of molecular evolutionary rates

Genetic polymorphism within a species is generated by past mutations that have neither reached fixation nor been deleted. The probability *q* that a site is polymorphic is formulated as

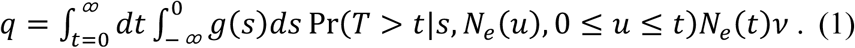

Here, *v* is the mutation rate, *s* and *g*(*s*) are the selection intensity and its distribution, and *N*_*e*_(*u*) is the effective population size at time *u* before present. We note that the probability of a positive value for *s* can be assumed to be negligible. *T*, which is the survival time of a mutation before either being fixed in the population or deleted, depends on the selection intensity and population history. Ignoring the contribution of slightly deleterious mutations for simplicity, we consider the scenario in which genetic polymorphism is largely generated by neutral mutations. In this scenario, (1) becomes

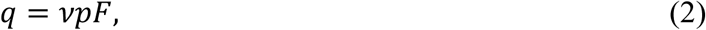

where

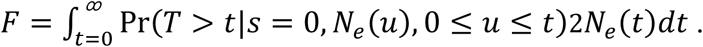

The probability that a mutation is selectively neutral, *p*, expresses the extent of the independence of the mutation from functional constraints. While *p* varies among genes, *F* does not. As a result, among-gene variation of the proportion of segregating sites reflects either the variation of mutation rates or the variation of functional constraints.

Interspecies molecular evolution, in contrast, is based on mutations that have become fixed in populations. Assuming the above scenario, the neutral theory of molecular evolution expresses the molecular evolutionary rate as

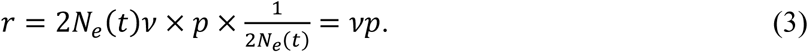

Although intraspecies genetic variation is many orders of magnitude lower than that generated during the evolution of all mammals, equations (2) and (3) predict that the among-gene variation of the proportion of segregating sites is correlated with the among-gene variation of the molecular evolutionary rate.

The latter rate may be measured from the variation in total branch lengths of gene trees, which expresses the number of substitutions across the whole tree. At the same time, mean branch lengths of gene trees should well reflect the variation in evolutionary rates among genes, even in the presence of missing values, unless the resultant sub-trees of the (unobserved) gene trees are seriously skewed. Gene trees sometimes include extremely long terminal branches, even after careful inspection to control the quality of the alignment. Because mean branch length is sensitive to extremely long outlier branches, we also calculated the median of branch lengths as a robust estimate of among-gene variation in molecular evolutionary rates.

### Neutral theory extended to multiple-gene molecular evolution

The well-accepted neutral theory of molecular evolution asserts that selectively advantageous mutations are rare among those mutations leading to interspecies variation [4]. As a result, the rate of molecular evolution is the product of the mutation rate, *v*, and the proportion of neutral mutations, *p*[6]. By extending this relationship to the rates of multiple-gene molecular evolution, we obtained the following formula for the branch lengths of gene trees [9]:

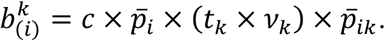

Here, 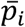 represents the relative variability of gene *i* (*i* = 1, …, *N*), and *t*_*k*_ and *v*_*k*_ are the evolutionary time span and mutation rate along branch *k* (*k* = 1, …, *M*). By applying a two-way ANOVA-type Poisson regression to the scaled branch lengths 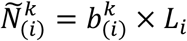 (where *L*_*i*_ is the length of gene *i*), namely,

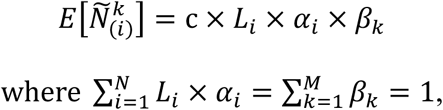

we can estimate 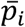 and *t*_*k*_ × *v*_*k*_ based on the gene effect 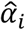 and branch effect 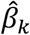. The interaction 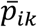, which captures the among-branch variation of functional constraints on a gene, is estimated as the ratio of 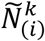 to the predicted value. When genomes include a complete set of orthologous genes for all species, the maximum likelihood estimators of the gene effect and the branch effect can be obtained as follows:

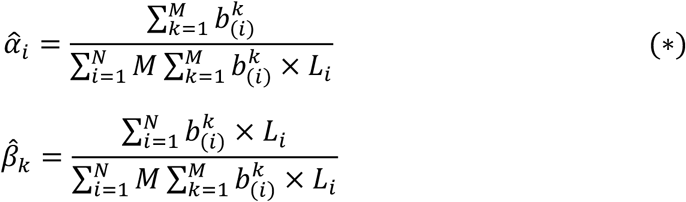

### Summary statistics describing variation in long-term evolutionary rates among genes in a genome

In this study, we investigated the possibility that intraspecies population genetic diversity is affected by the between-species long-term molecular evolutionary rate. Specifically, we contrasted the rate of segregation in humans with the rate of molecular evolution in mammals. As can be seen in equation (*), the among-gene variation of the molecular evolutionary rate can be measured from the total branch lengths of gene trees.

In actual practice, however, gene trees often include missing taxa, partly because of incomplete sequencing but mainly because of gene loss over the course of mammalian evolution. As a result, the among-gene variation of total branch lengths reflects not only the among-gene variation of molecular evolutionary rate but also variation in taxon sampling. To reduce the effect of differential taxon sampling, we used mean branch lengths:

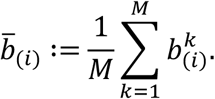

The variation in mean branch length should well reflect the variation in evolutionary rates among genes, even in the presence of missing values, unless the resultant sub-trees of the (unobserved) complete gene trees are seriously skewed.

The generated gene trees sometimes included extremely long terminal branches. Long branches in a gene tree may be ascribed to three phenomena: 1) sequencing or annotation errors, 2) bias caused by the model used for tree inference, or 3) lineage-specific selection. Even after careful inspection to control the alignment quality, however, we could not exclude the possibility of incomplete modeling of molecular evolution or lineage-specific selection in some cases. Because mean branch length is sensitive to extremely long outlier branches, we also calculated the median of the branch lengths as a robust estimate of the among-gene variation in molecular evolutionary rates:

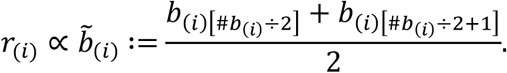

With regard to protein evolution at the amino acid level, more than half of the branches of some extremely conserved genes were of zero length; this resulted in a median branch length of 0, which would be problematic in the succeeding regression analysis. We therefore removed the five longest branches from each gene tree and obtained the truncated mean of branches, 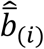, as an estimate of 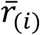. Both 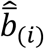 and 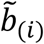 were used in this study.

### Observed and corrected correlations between intraspecies genetic diversity and long-term molecular evolutionary rates

To examine the effects of long-term molecular evolutionary rates on intraspecies genetic diversity, we calculated the correlation between the rate of molecular evolution *B* = {*b*_(1)_, *b*_(2)_, …, *b*_(*i*)_, …} and the number of segregating sites *K* = {*K*_(1)_, *K*_(2)_, …, *K*_(*i*)_, …}. Because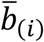 and *K*_(*i*)_ were accompanied by the random noise of estimation and sampling, we corrected the observed correlation by taking into account the sampling variance of each 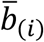 and *K*_(*i*)_.

To investigate the effect of sampling variance, we denote the observed values of *B* and *K* as 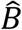 and 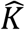, where 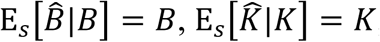, and the subscript “S” represents the sampling mean and variance. In addition, we use the subscript “G” to represent the among-gene mean and variance. To the first order, the expectation of the observed correlation between 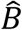 and 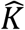,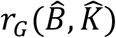, is contrasted with *r*:(*B, K*) as follows:

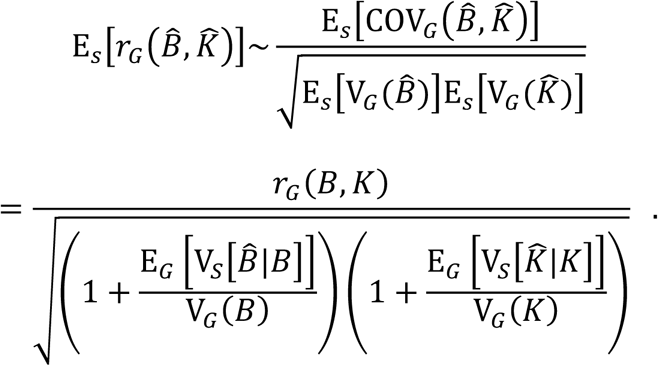

By evaluating the sampling variances of 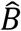 and 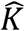, we obtain the bias-corrected correlation as follows:

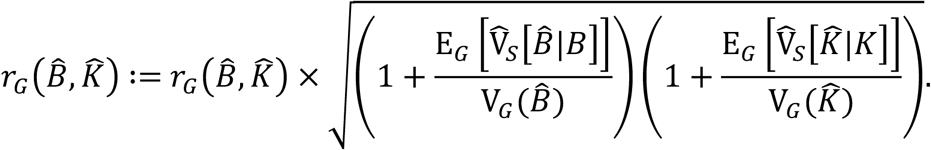

### Sampling variance of mean branch lengths

We evaluated the sampling variance of branch lengths with reference to the Poisson random variables for the numbers of nucleotide or amino acid substitutions. Given that the branch length of gene *i* of branch, 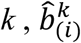, is the expected number of substitutions per site, we assume that the stochastic variance of 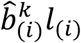 is approximated by the variance of a Poisson random variable with mean 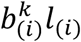, where *l*_(*i*)_ is the sequence length of gene *i*:

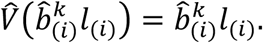

The variance of the mean branch lengths of gene *i* is therefore evaluated as

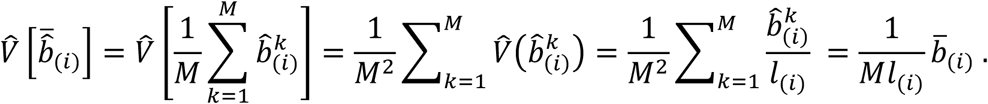

Bootstrap resampling of columns in the alignment would allow estimation of the variance of median and mean branch lengths. As an alternative approach, we developed a simple simulation method to evaluate the stochastic variance due to the randomness of substitutions. Given a gene tree, we generated the number of substitutions for each branch from the Poisson distribution, with ‘its mean set to the product of the branch length and the gene length. We obtained the sample of the mean (and median) branch length by dividing the mean (and median) of the generated number of substitutions by the gene length.

### Stochastic variance of intraspecies genetic diversity

If one assumes that mutations in a population are neutral and that the population is at equilibrium, the variance in the number of segregating sites, *K*, is

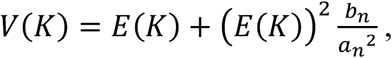

where 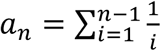 and 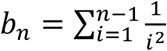. Here, *n* is the number of chromosomes in a sample from the population [31]. The first term reflects the randomness of mutations, and the second term measures the stochasticity of the genealogy. When the population is not at equilibrium but has its own history, the variance becomes

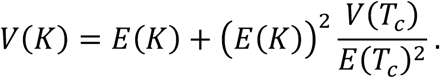

Here, *E*(*T*_c_) and *V*(*T*_c_) are the mean and variance of waiting times of coalescence events in genealogies [15]. Unfortunately, detailed information on genealogies is not available in most cases. We note, however, that the above variance formula is consistent with the variance formula of a negative binomial distribution. We therefore applied negative-binomial regression with a log link:

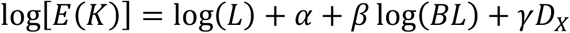

where *L* = {*l*_(1)_, *l*_(2)_, …, *l*_(*i*)_, …} is the length of genes and *BL* is the mean branch length. *D*_X_ is a dummy variable for genes on the X chromosome that incorporates differences in effective population size between an autosome and the X chromosome. We carried out this regression using the function glm.nb in the R package MASS [32]. The variance of *K* can be obtained by

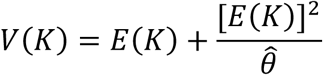

where 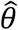 is the shape parameter of the negative binomial distribution.

We also applied the model

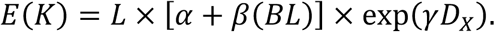

Maximum likelihood inference was conducted by applying the function optim. A positive value of *α* implies the intraspecies survival of mutations destined for deletion from the population on a macroevolutionary time scale.

### Species specificity: the significant deviation of trait-associated gene sets

Intraspecies genetic diversity is affected by long-term molecular evolutionary rates. Species-specific enhanced/reduced functional constraints may be observed as residuals of the prediction of the above negative binomial regression. For each gene in a trait-associated gene set, we calculated the *p*-value of the observed number of segregating sites based on the predicted value and the estimated scale parameter. To obtain the significance of the gene set, we standardized the *p*-values by transforming them to the percentiles of their corresponding standard normal distribution. The *p*-value of the sample mean of these transformed values was obtained using a *t*-distribution as a reference. The *t*-values measure the deviation from the predicted genetic diversities of genes in gene sets. Positive *t*-values imply the existence of reduced functional constraints or diversifying selection in a human lineage, whereas negative values indicate the presence of enhanced functional constraints or purifying selection. Given the false discovery rate value, significant genes were selected by the Benjamini and Hochberg procedure [33].

### Multiple alignment of mammalian genes

We downloaded 96 complete mammalian genomes from GenBank and used a custom Perl script to extract protein-coding sequences of each species. A gene pool containing 21,350 mammalian genes was generated based on NCBI genomic annotation. We generated a multi-sequence file of all genes in the gene pool, and kept one sequence per species for each gene. Including genes that are shared by more than 70 species out of 96 mammals, and also with the human 1000 Genomes Project annotation (hg19 human genome assembly), a total of 14, 671 genes were used in this study. Among these genes, 5,560 were previously reported as single-copy genes in Class Mammalian in Wu et al. (2017) [9]. Alignments at the codon level were performed in Prank v.170427 [34]. Sites with less than 70% coverage among all species, as well as sequences with less than 30% coverage among gene loci in the alignment, were removed.

### Inference of gene trees

We generated a maximum likelihood tree for each gene using IQ-TREE software [35], which automatically performed model selection and determined the best data partitions. The best evolutionary model for each gene was independently selected based on the Bayesian information criterion and used to infer the nucleotide tree. All gene trees were calculated using 1,000 bootstrap replicates.

### Population data

We downloaded single-nucleotide polymorphism (SNP) data for 2,504 humans (*Homo sapiens*) from the 1000 Genomes Project (Phase 3 data) [10-12]. SNP data for 60 common chimpanzees (*Pan troglodytes*), 31 gorillas (*Gorilla gorilla*), 35 laboratory mice (*Mus musculus*) and 127 dogs and wolves (*Canis lupus*) were collected from published papers and databases [36-39]. Information on *de novo* mutations in the human genome was collected from Jonsson et al. [1]. Human ancestral alleles were collected from Keightley and Jackson [40]. Individual-derived alleles of each gene were collected by comparing SNPs of each individual with human ancestral sequences.

## Data availability

Source data and gene trees are available at https://github.com/wujiaqi06/Long-Term-Rate.

## Acknowledgements

This study was supported by a Grant-in-Aid for Scientific Research (B) (19H04070) and a Grant-in-Aid for JSPS Research Fellows (18F18385) from the Japan Society for the Promotion of Science. We deeply thank Jeffrey L. Thorne of North Carolina State University, Zhenqiu Liu of Fudan University and Nikaido Masato of the Tokyo Institute of Technology for their valuable comments and suggestions on our manuscript. We thank Barbara Goodson, from Edanz Group (www.edanzediting.com/ac) and Ian Smith from Elite Scientific Editing (http://www.elitescientificediting.co.uk), for editing the English text of a draft of this manuscript.

## Contributions

J.W., T.Y., and H.K. designed the research. J.W. and H.K. formulated the methods and performed the data analysis. J.W., T.Y., and H.K. wrote the paper.

## Correspondence and requests for materials

Correspondence and requests for materials should be addressed to http://wujiaqi06@gmail.com

## Supplemental table captions

**Figure S1. Time scales of mammalian and human evolution. A**. The differing evolutionary time scales of Class Mammalia, Subfamily Homininae, and *Homo sapiens*. The time tree for mammals is from dos Reis (2012) [13]. **B**. Examples of mammalian gene trees. **C**. Examples of population-level genetic diversity, including indels and single-nucleotide polymorphisms (SNPs).

**Figure S2**. Correlation between the proportion of segregating sites (*q*) and the rate of mammalian molecular evolution (*r*) of chimpanzee, gorilla, mouse and dog/wolf. The result was obtained from 12,236 genes that are shared by all five species.

**Table S1. Top 20 genes for each principal component (PC) from PC1 to PC3 in principal component analysis**. The results of gene enrichment analysis are also summarized in this table.

**Table S2. Significant genes detected by correspondence analysis with a false discovery rate of 0**.**01**. The results of gene enrichment analysis are also summarized in this table.

**Table S3. *p*-values and *t*-values of each disease in disease-associated gene set tests of all 14,267 diseases of human, chimpanzee, gorilla, mouse and dog/wolf**.

**Table S4. Significant disease-associated gene sets in human, chimpanzee, gorilla, mouse and dog/wolf, with a false discovery rate of 0.01**.

**Table S5. Significant disease-associated gene sets detected by correspondence analysis using −*log*_10_(*p*) of each disease-associated gene set, with a false discovery rate of 0.01**.

**Data S1. Gene tree of 96 mammalian species for 21, 350 genes**.

**Data S2. Mammalian long term rates, proportion of segregating sites (*q*), proportion of *de novo* mutations, proportion of singletons, nucleotide diversity, etc. of human**.

**Data S3. Mammalian long term rates and proportion of segregating sites (*q*) of five species**.

