## Supplementary figures and images for "Long-term molecular evolutionary rate determines intraspecific genetic diversity"

### Figure 1

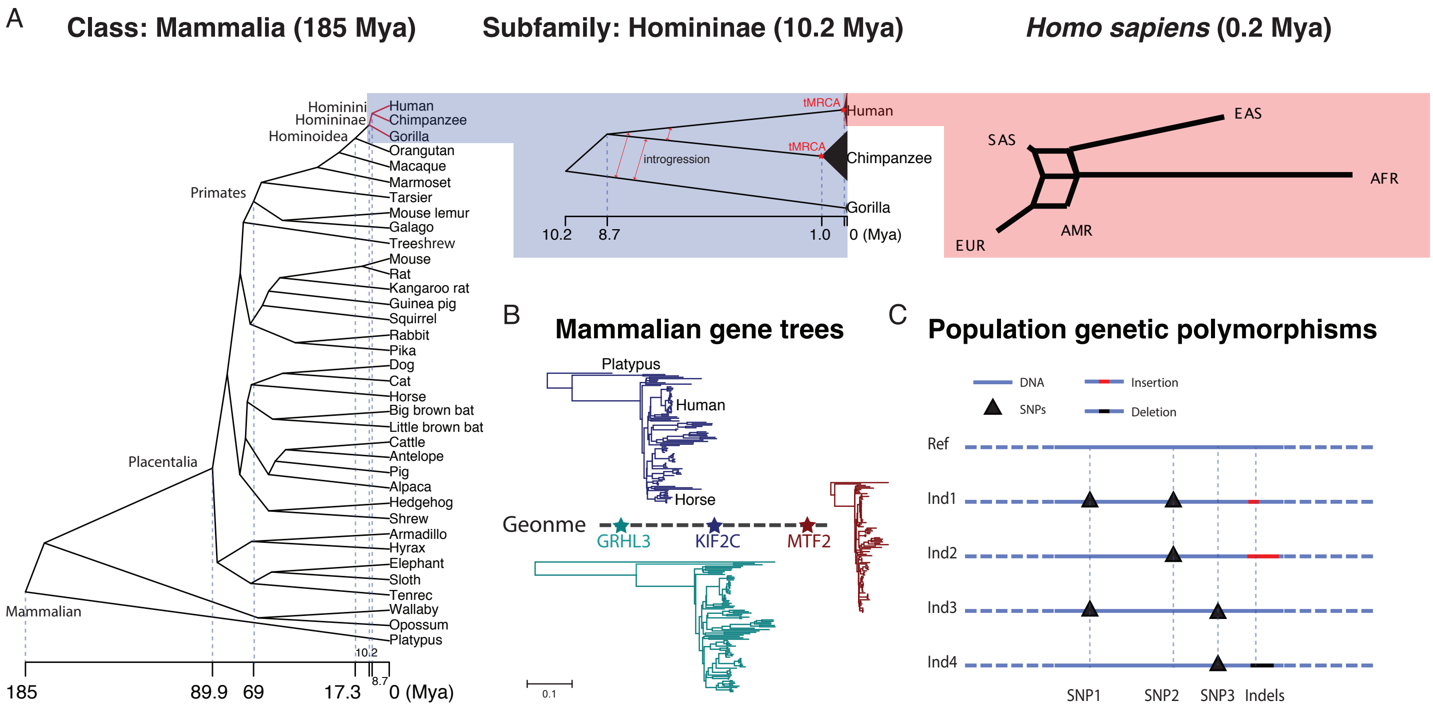

### Figure 1

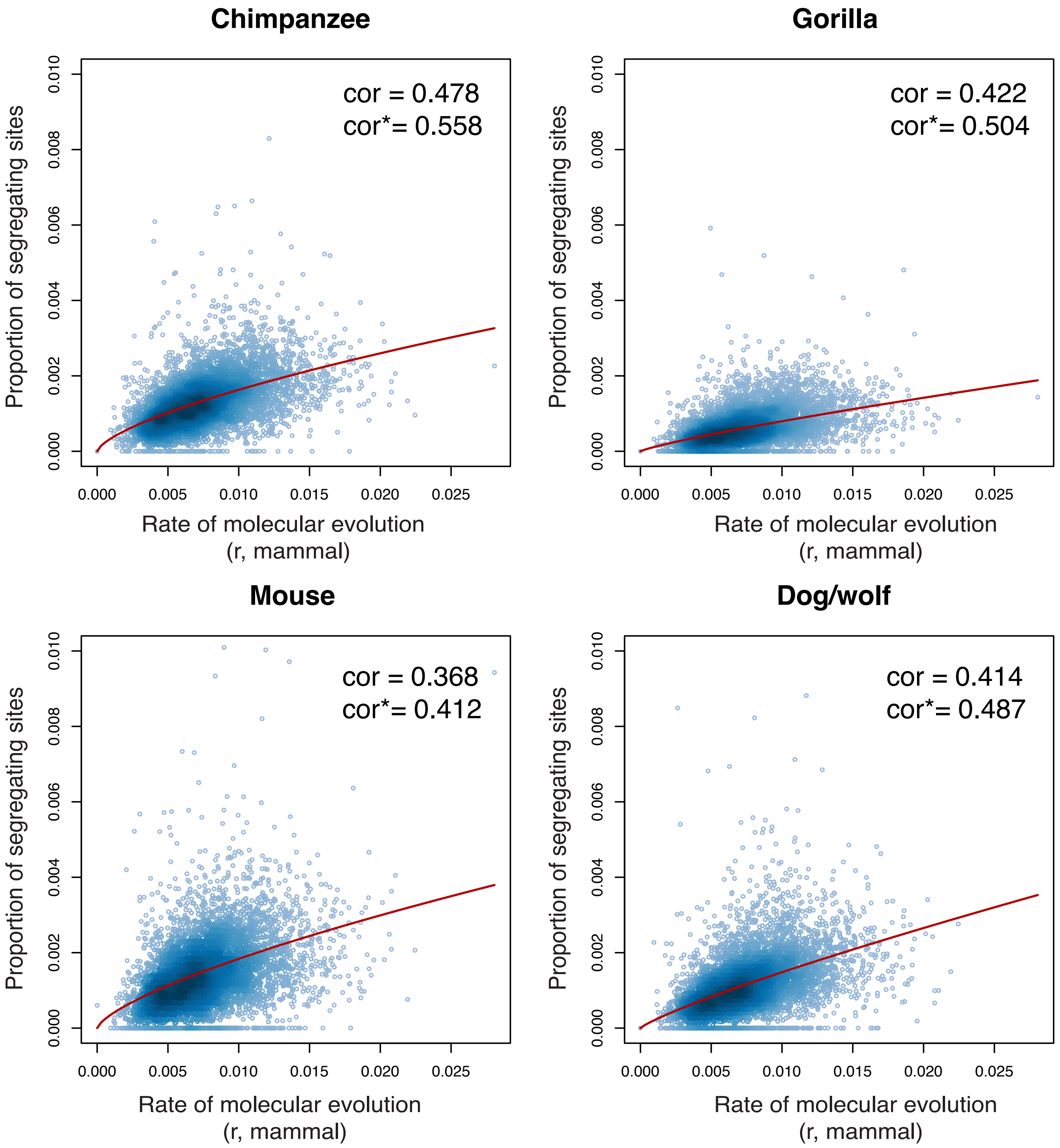
